# ComI inhibits transformation in *Bacillus subtilis* by selectively killing competent cells

**DOI:** 10.1101/2024.02.02.578676

**Authors:** Dominique R. Smith, Daniel B. Kearns, Briana M. Burton

**Affiliations:** Department of Bacteriology, University of Wisconsin – Madison, Madison, WI, USA; Department of Biology, Indiana University, Bloomington, IN, USA

**Keywords:** competence, natural transformation, *B*. *subtilis*

## Abstract

Many bacteria build elaborate molecular machines to import DNA via natural competence, yet this activity is often not identified until strains have been handled and domesticated in laboratory settings. For example, one of the best studied Gram-positive model organisms, *Bacillus subtilis,* has a non-transformable ancestor. Transformation in the ancestral strain is inhibited by a transmembrane peptide, ComI, which is encoded on an extrachromosomal plasmid. Although ComI was shown to be necessary and sufficient to inhibit transformation when produced at high levels under an inducible promoter, the mechanism by which ComI inhibits transformation is unknown. Here, we examine the native regulation and mechanism of transformation inhibition by ComI. We find that under native regulation, ComI expression is restricted in the absence of the plasmid. In the presence of the plasmid, we find that ComI is preferentially expressed in cells that are differentiating into a competent state. The subcellular localization of ComI, however, does not depend on any other competence proteins and permeabilization activity is concentration dependent. Thus over time, the competent cells gradually producing ComI, are permeabilized and killed. Based on these observations we propose a new model for the mechanism of ComI, suggesting a response to competence activation that selectively eliminates the competent subpopulation.

**Importance:** Natural transformation mechanisms have been studied across several bacterial systems, but few examples of inhibition exist. This work investigates the mechanism of action of a plasmid-encoded transmembrane inhibitor of natural transformation. The data reveal that the peptide can cause cell permeabilization. Permeabilization is synergistic with entry of *Bacillus subtilis* into the “competent” state, such that cells with ability to be transformed are preferentially killed. These findings reveal a self-preservation mechanism coupled to the physiological state of the cells that ensures the population can maintain unaltered plasmid and its predicted prophage.

## Introduction

Natural competence refers to the physiological state that enables bacteria to take up genetic material from their environment (1). A complex protein apparatus is assembled at the cytoplasmic membrane and is required for the uptake of DNA into the cytoplasm. Upon entry into the cell, single-stranded DNA (ssDNA) can serve as a substrate for chromosomal DNA repair or plays a role in the acquisition or deletion of genes (2, 3). Natural competence is an important factor in the evolution of bacteria, contributing to the emergence of pathogens, the dissemination of virulence factors, and mediating the spread of antibiotic resistance genes (1, 4). In *B. subtilis,* a competence-specific regulon is activated by the master regulator ComK (5). In some species, despite the presence of complete regulons for natural competence, natural transformation has not been demonstrated. In fact, the genes for competence are common, but the procurement of natural transformants is relatively rare. One reason for this may be the presence of transformation inhibitors.

Domestication made *B. subtilis* a model organism due to its ability to grow rapidly and integrate DNA into its genome, but the ancestor from which lab strains were derived (NCIB3610, hereafter referred to as 3610), was poorly competent (6). Competence was inhibited in 3610 by the presence of the plasmid pBS32 and cured strains were nearly as transformable as the laboratory strains (7–10). Nested deletion analysis of pBS32 mapped the inhibitory activity to a gene called *comI*, which encodes a 30 amino acid predicted single-pass transmembrane alpha helix (10, 11). ComI was shown to be sufficient to inhibit transformation because a strain lacking the plasmid and ectopically expressing inducible *comI* on the chromosome inhibited transformability 100-fold or more (10). During domestication of *B. subtilis*, the plasmid was naturally cured and ComI was lost. The mechanism of ComI-mediated competence inhibition is poorly understood and the biological reason that a horizontally transferred element like a plasmid encodes an inhibitor of horizontal gene transfer is unknown.

One model of transformation inhibition argues that ComI interacts with and somehow disrupts part of the DNA uptake machinery. ComI appears to inhibit transformation post-transcriptionally because the frequency of gene expression of a transcriptional fusion driven by the *comK* promoter in 3610 was only 10-fold lower than in PY79, and it was suggested that this would not account for the 1000-fold difference in transformation frequency between the strains (10) Moreover, ComI appears to be a transmembrane helix, and a glutamine-to-leucine substitution in the transmembrane residue glutamine 12 significantly restores the transformation frequency, perhaps suggesting specificity-of-interaction within the membrane bilayer (10). Finally, fluorescent puncta of an inducible mCherry-ComI fusion co-localized with puncta of GFP fused to a late competence protein, ComGA (10, 12). Altogether, these data led to the proposal that ComI could be interacting with one of the transmembrane competence machinery proteins via glutamine 12 (10, 13, 14). Whether ComI inhibits competence by direct interaction with the competence machinery is speculative as no direct interaction partner has been determined.

To date, ComI has been studied under an inducible promoter in a background lacking pBS32. It was demonstrated that transformability decreased significantly at final induction concentrations of ≥ 70 μM IPTG, while at ≥ 80 μM IPTG induction, cell viability decreased considerably (10). It was speculated that the drop in cell viability was the result of membrane toxicity from artificially overexpressed ComI that did not occur in the wild type (10). Here we investigated the tight correlation between loss of transformability and viability. We investigated the possibility that when under native control, ComI might be produced at higher levels during competence and that transformation inhibition could be the direct result of ComI-mediated cell death. Our results show that inducible ComI permeabilizes the cells in a concentration dependent manner. Cells cultured under competence conditions show greater growth inhibition by ComI, in comparison to cells cultured in non-competence inducing conditions. Further, we determine that only the competent subpopulation of cells has significantly increased transcription from the *comI* promoter and that these cells die. Together our data demonstrate that ComI selectively permeabilizes the competent subpopulation leading to cell death.

## Results

### Inducible ComI Permeabilizes the Membrane Resulting in Cell Death

It was previously shown that during a transformation experiment, a strain with inducible ComI had a sharp drop in viability when the inducer exceeded a final concentration of 70 μM. To understand the toxicity of ComI when under IPTG induction, we analyzed the growth behavior of PY79 cells expressing low or high levels of ComI in non-competent (LB) and competent (MC) growth conditions. ComI was integrated at the chromosomal locus *amyE,* and gene expression was driven by the IPTG-inducible *hyperspank* promoter. Inducer was added either directly to cells at the point of dilution from an overnight culture or to cells that were in the exponential growth phase. In the first scenario, when cells from an overnight culture were inoculated into fresh medium and simultaneously induced with 10 μM IPTG, the growth was nearly identical to uninduced cells (Figure 1A, 1B). When the final concentration of inducer was raised to 70 μM, cell growth was slightly reduced. By contrast, 100 μM IPTG induction prevented reinitiation of growth (Figure 1A, 1B). Titration suggested that there was a critical induction concentration required to promote a toxic effect on the cells independent of the growth conditions. In the second scenario, inducer was added to cells that were in exponential growth and a distinct growth pattern was observed. Cells induced with 70 and 100 μM IPTG completed one or two rounds of doubling before growth rate plateaued and optical density even began to decrease (Figure 1C, 1D). The effect of ComI induction during exponential growth was more pronounced in MC compared to cells growing in LB. In particular, expression was pronounced at the onset of the optimal competence window, indicating that cells grown in competence conditions were more susceptible to ComI toxicity (Figure 1C, 1D). To confirm that the IPTG was not having an indirect effect on cell growth, we also followed growth of cells with an empty *hyperspank* promoter at the same ectopic site in the chromosome. These cells showed no impairment in growth in either condition (Figure S1).

**Figure 1.**
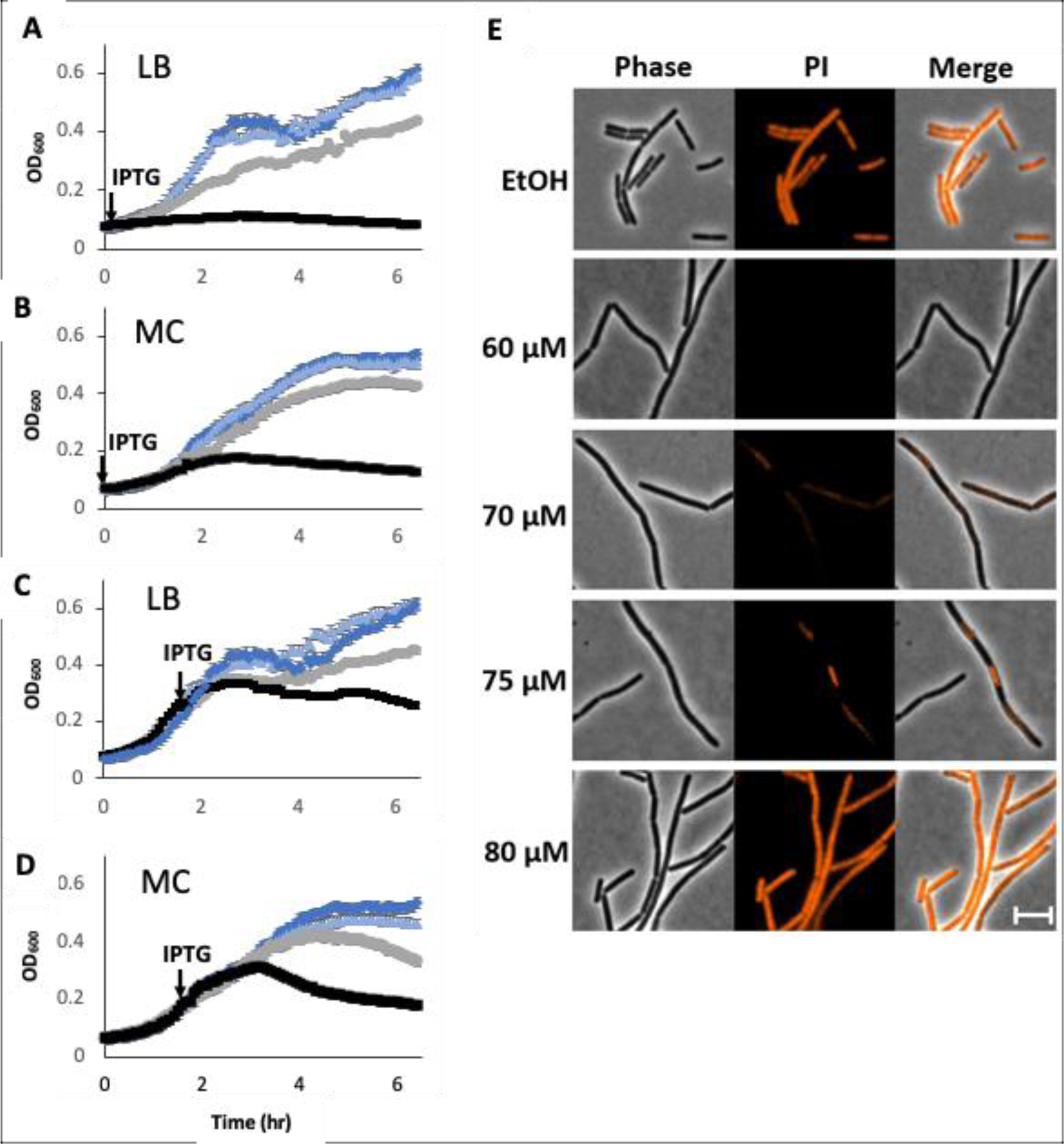
Inducible ComI permeabilizes cells resulting in growth impairment. A and C) Growth of cells in LB that were induced at the time of inoculation or during exponential phase growth respectively. B and D) Growth of cells in MC that were induced during pre-logarithmic and logarithmic growth respectively. Dark blue 0 μM, light blue 10 μM, light grey 70 μM, black 100 μM. Errors bars represent the standard deviations of 3 replicates. E) Inducible ComI stained with 300 ng/uL propidium iodine (false colored orange). Cells were grown in non-competence conditions and were induced 1.5 hours pre-imaging. The white scale bar on the bottom panel represents a distance of five microns. A PY79 background containing *amyE::P_hyperspank_-comI* (bBB592) was used to generate panels A-E.

Next, we investigated if the observed growth defects stemmed from cell permeabilization. We used the same conditions that produced ComI toxicity and then subjected the cells to propidium iodide (PI). Propidium iodide cannot enter cells with an intact cell membrane; thus, if production of ComI disrupts membrane integrity, PI can traverse the membrane, intercalate in the DNA and result in fluorescent cells (15). Using the same inducible strain as above, IPTG was titrated, and we examined cells for PI signal above background. Induction of ComI at up to 60 μM IPTG resulted in no increased PI signal above background (Figure 1E). Induction of ComI at ≥ 70 μM IPTG for 1.5 hours resulted in elevated PI signal in a subpopulation of the cells. The PI signal was the highest and present in most of the cells at 80 μM IPTG induction (Figure 1E). The increase in both the fraction of cells with signal above background and total PI signal intensity per cell indicates that the cells became permeable at a critical concentration.

The amount of inducer required for cell permeabilization aligns closely with a previously reported IPTG titration which indicated a significant decline in cell viability starting at 80 μM IPTG (10). We began to detect an increasing number of permeable cells starting from 70 μM IPTG induction (Figure 1E). This raised the possibility that viability may be compromised at a lower concentration of inducer, one that was previously reported to only lead to a reduction in transformation frequency (10). Considering these data, we next explored whether the behavior of inducible ComI depends on the presence of other components of the competence machinery.

### ComI Localizes as Discrete Puncta at the Membrane Independent of the Competence Machinery

It has been observed that ComI may colocalize with the competence machinery (10). Although the Com-dependent permeabilization we observed could happen in non-competence conditions, we wanted to investigate the potential connection between ComI and competence machinery further. To do this, we first compared the localization patterns of mCherry-ComI in competent and non-competent conditions. To ensure the competence machinery did not form in our non-competent condition, we expressed mCherry-ComI in a strain that had a deletion of the chromosomally encoded *comK,* the master regulator of competence (Figure 2). In the *comK* deletion strain, no late competence proteins should be made even when cells are grown in competence promoting MC medium. Therefore, if mCherry-ComI puncta localized due to association with one of the Com proteins, this localization pattern might change in the *comK* deletion strain. When mCherry-ComI was ectopically expressed in a 3610 cured background that had a deletion in the chromosomally encoded *comK*, we observed a similar pattern of discrete puncta distributed around the cell as that seen in cells producing late competence machinery proteins (Figure 2). Thus, the localization pattern of ComI appears independent of the late competence proteins and further supports the possibility that ComI function does not require direct interaction with the competence machinery. Consistent with the results here, in the original colocalization studies, numerous cells with visible mCherry-ComI puncta did not have detectable late competence machinery puncta (10). To this point, the localization patterns and growth impairment could be the result of the overexpression of ComI. Therefore, to gain more physiologically relevant insight on ComI function, we investigated ComI under its native promoter.

**Figure 2.**
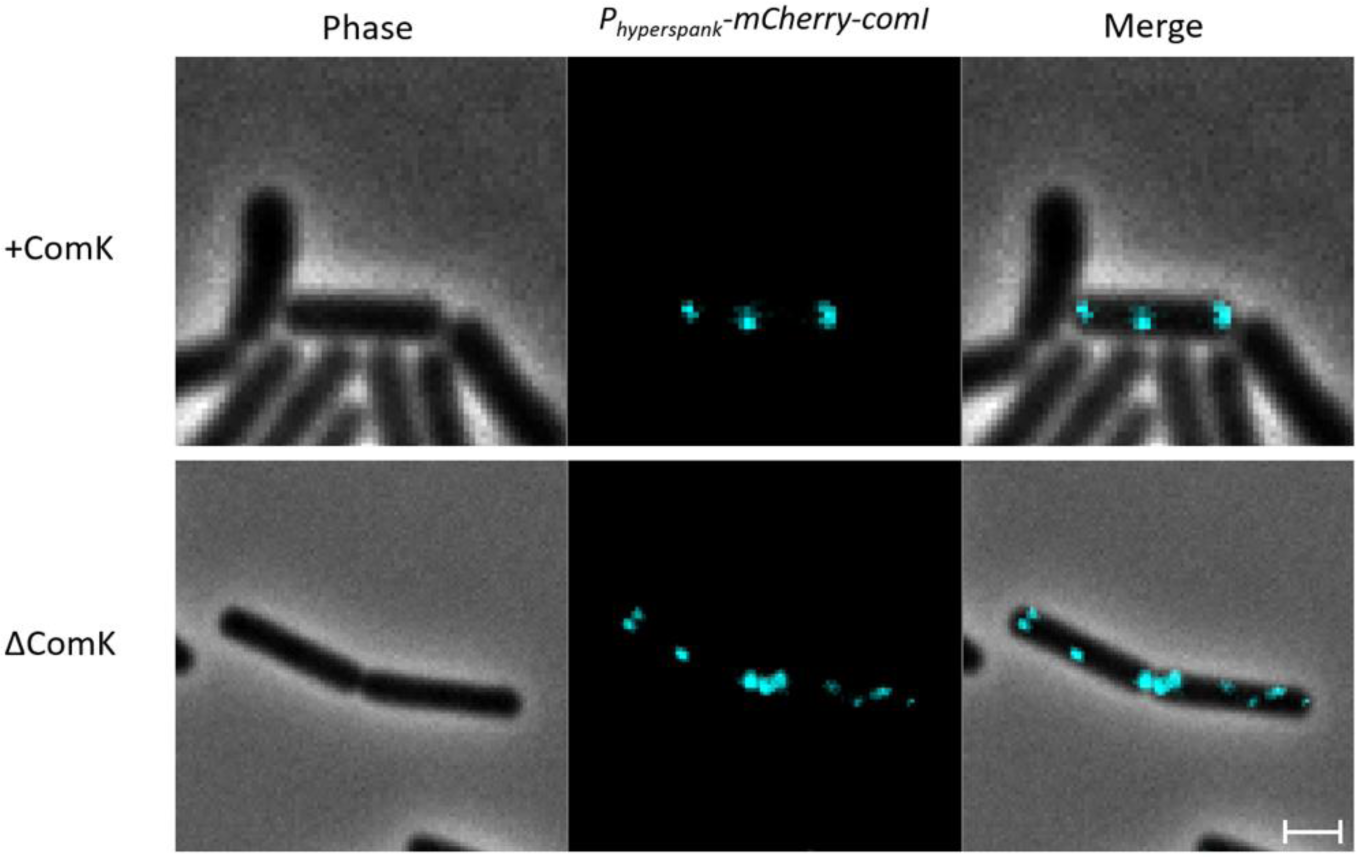
ComI localization is independent of competence machinery. A) Discrete mCherry-ComI puncta (false colored turquoise) are observed at the membranes. Micrographs of cells producing mCherry-ComI grown in MC and LB in the presence or absence of ComK respectively and were induced with 70 μM IPTG 1.5 hours pre-imaging. The white scale bar on the bottom panel represents a distance of one micron. Strains included in this figure are: bDS001 and bDS014.

### pBS32-Regulated ComI Inhibits Transformability but Does Not Decrease Bulk Cell Viability

To explore the intrinsic regulation of ComI, we first examined whether *comI* driven by its native promoter was sufficient for ComI-mediated inhibition. We integrated *comI* driven by its endogenous promoter into the chromosome in various genetic backgrounds (Figure 3A). We then determined transformation frequencies and viability for these strains. As previously demonstrated, wild-type 3610 had an approximate transformation frequency of 10^-6^ and curing the plasmid resulted in a 100-fold increase in transformation frequency (Figure 3B, black and dark blue) (10, 16). When *comI* was integrated on the chromosome in the cured strain, the transformation frequency was similar to the cured strain alone (Figure 3B, dark and light blue). We infer that in the absence of pBS32, no element from the chromosome directly upregulated the chromosomally encoded *comI*. We then asked whether an element on the plasmid was needed for ComI-mediated transformation inhibition. To answer this, we integrated *comI* on the chromosome in a background that had an in-frame deletion of *comI* on the plasmid. Indeed, when ComI was expressed under native control in the presence of the plasmid with the in-frame *comI* deletion, there was a 400-fold decrease in transformation frequency relative to the *ΔcomI* strain alone (Figure 3B, dark and light orange). These results suggested that when the plasmid is present, *comI* encoded in *trans* on the chromosome is produced at levels that are sufficient to inhibit transformation. Similarly, integrating *comI* on the chromosome in a strain harboring pBS32 with *comI* coding for the missense mutation Q12L resulted in a 200-fold decrease in the transformation frequency relative to the strain containing a Q12L mutation alone (Figure 3B, dark and light green). In this case, although a non-functional copy of ComI is being made on the plasmid, the chromosomally encoded ComI still significantly inhibits transformability. Combined, these data established that native levels of ComI are sufficient to inhibit transformation in a pBS32-dependent manner.

**Figure 3.**
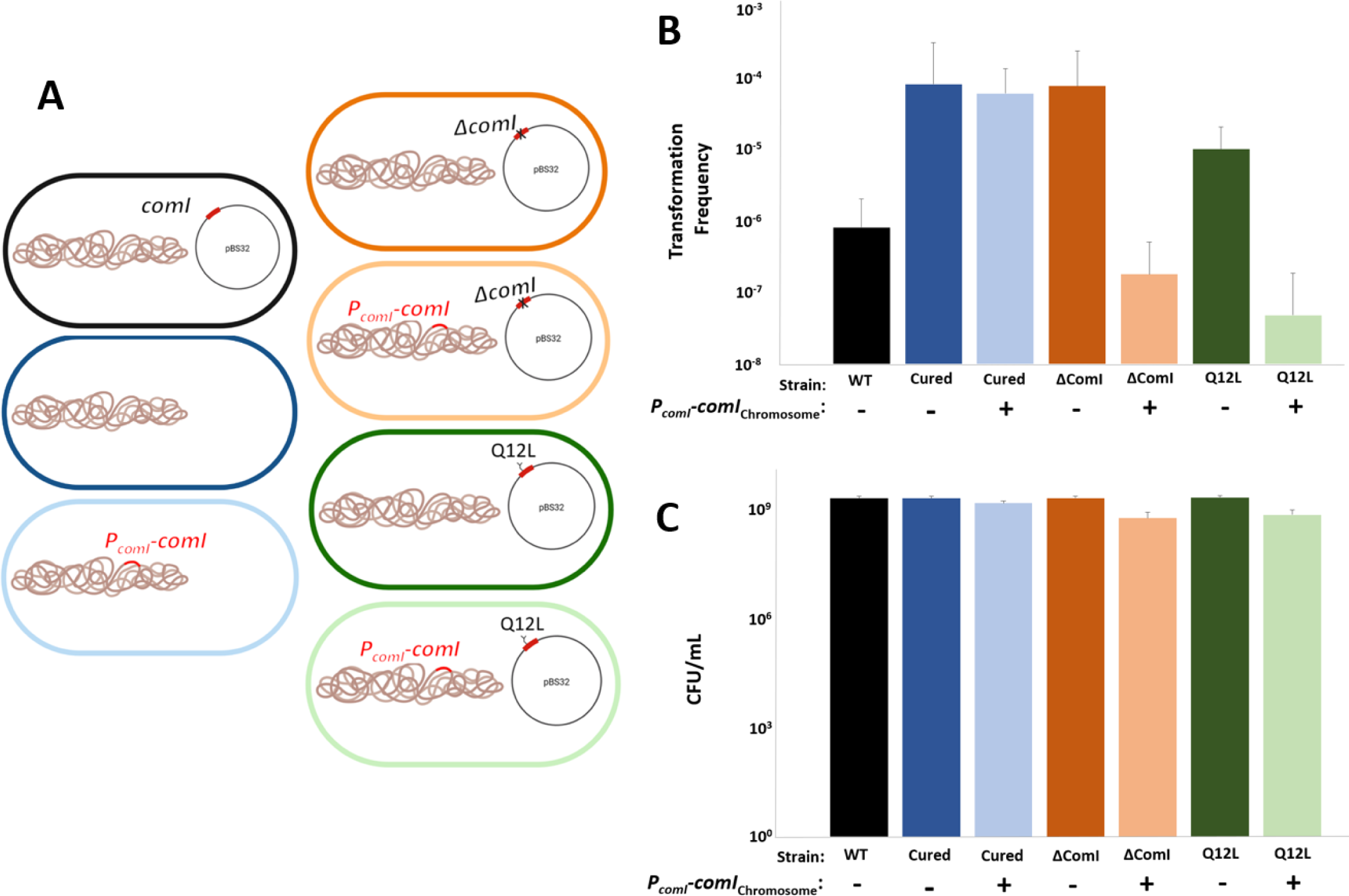
Transformation inhibition by natively expressed ComI is Dependent on pBS32 and does not decrease cell viability. A) Schematic representation of each strain genotype. The strains are color-coded to match the transformation frequency and the cell viability bar graphs. B) Transformation frequencies of WT and mutant genotypes. C) Cell viability associated with each transformation frequency. Graphs contain error bars that are the standard deviations of three replicates. Strains are described on the x-axis and *P_comI_-comI*_chromosome_ indicates the presence of natively expressed ComI at the chromosomal locus *ytoI.* The following 3610 strains were used in this figure: WT (bLH080), cured (bBS108), *ytoI::P_comI_-comI* (bDS005), *ΔcomI* (bBB592), *ytoI::P_comI_-comI ΔcomI* (bDS004), ComI Q12L (bJF176), *ytoI::P_comI_-comI* ComI Q12L (bDS003).

Despite observing a direct correlation between the loss in transformability and cell viability when ComI was expressed under the inducible promoter, no correlation was observed between these factors when ComI was expressed from its native promoter (Figure 3C). There was no significant difference in total cell viability on agar plates between strains that inhibited or did not inhibit transformation frequency (Figure 3C). We postulated that if the natively regulated ComI does not result in a global decrease in cell viability, perhaps it could preferentially kill a subpopulation of the cells. Since competence develops in a small proportion of cells, we wondered if ComI activity preferentially impacts the competent (hereafter referred to as *comK*-ON) population (17–19).

### *comK*-ON Cells have Significantly Increased *comI* Reporter Levels

To explore whether ComI could be preferentially acting in *comK-*ON cells, we turned to fluorescence microscopy. We built a dual-reporter system with transcriptional fusions of the *comI* and *comK* promoters to fluorescent proteins and integrated them at different ectopic loci in the chromosome (Figure 4A). In the cured strain, typically 10-20% of cells become *comK*-ON, and signal is detected from the *comK* reporter as early as 3.5 hours of growth in competence conditions. Signal from the *comI* reporter is detectable in all cells even as early as 2 hours regardless of growth conditions (not shown), and signal intensity does not change in the cells even at 4 hours of growth in *comK*-OFF cells, suggesting that *comI* is constitutively expressed (Figure 4). This is consistent with RNA-seq data which mapped ComI transcripts at 2 hours of growth in non-competence conditions (20, 21). However, to understand if there was a correlation in expression from the *comK* and *comI* promoters, we quantified the differences in *comI* reporter levels between the *comK*-ON and -OFF cells. In the cured background, *comK*-ON and -OFF cells had similar levels of GFP signal driven by the *comI* promoter (Figure 4B, 4E). However, in the presence of pBS32, we observed that *comK*-ON cells had significantly increased GFP signal from the *comI* promoter relative to *comK*-OFF cells (Figure 4C, 4F). Additionally, in the presence of pBS32, *comK*-ON cells were no longer phase-dark and appeared translucent, which suggests the cells were dead (Figure 4C). Although we observed drastic increases in *comI* reporter levels in WT 3610 cells, we could not calculate statistically significant differences in *comI* reporter levels between *comK*-ON and *comK*-OFF cells because only approximately 0.1% of cells were *comK*-ON at 5.5 hours of growth in competence conditions (Figure S3, 4C). If the loss of viable *comK*-ON cells is attributable to ComI, then deletion of c*omI* from pBS32 should increase throughput of *comK*-ON cells and will allow for quantification of *comI* reporter levels in *comK*-ON and -OFF cells. Considering this, we integrated the fluorescent reporter constructs into the strain containing the in-frame deletion of *comI* on pBS32. In this background, the frequency of cells that were *comK*-ON increased approximately 30-fold at 5.5 hours of growth in competence conditions (Figure 4D and S3). The *comI* reporter levels increased significantly in *comK-*ON cells relative to *comK*-OFF cells in the Δ*comI* background, and these cells remained phase-dark throughout growth (Figure 4D, 4G). Altogether, these data indicate that the presence of pBS32 is required for transcriptional activation of *comI*, that this upregulation is specific to cells that have differentiated into the *comK*-ON state, and that the viability of *comK*-ON cells is affected in a ComI-dependent manner.

**Figure 4.**
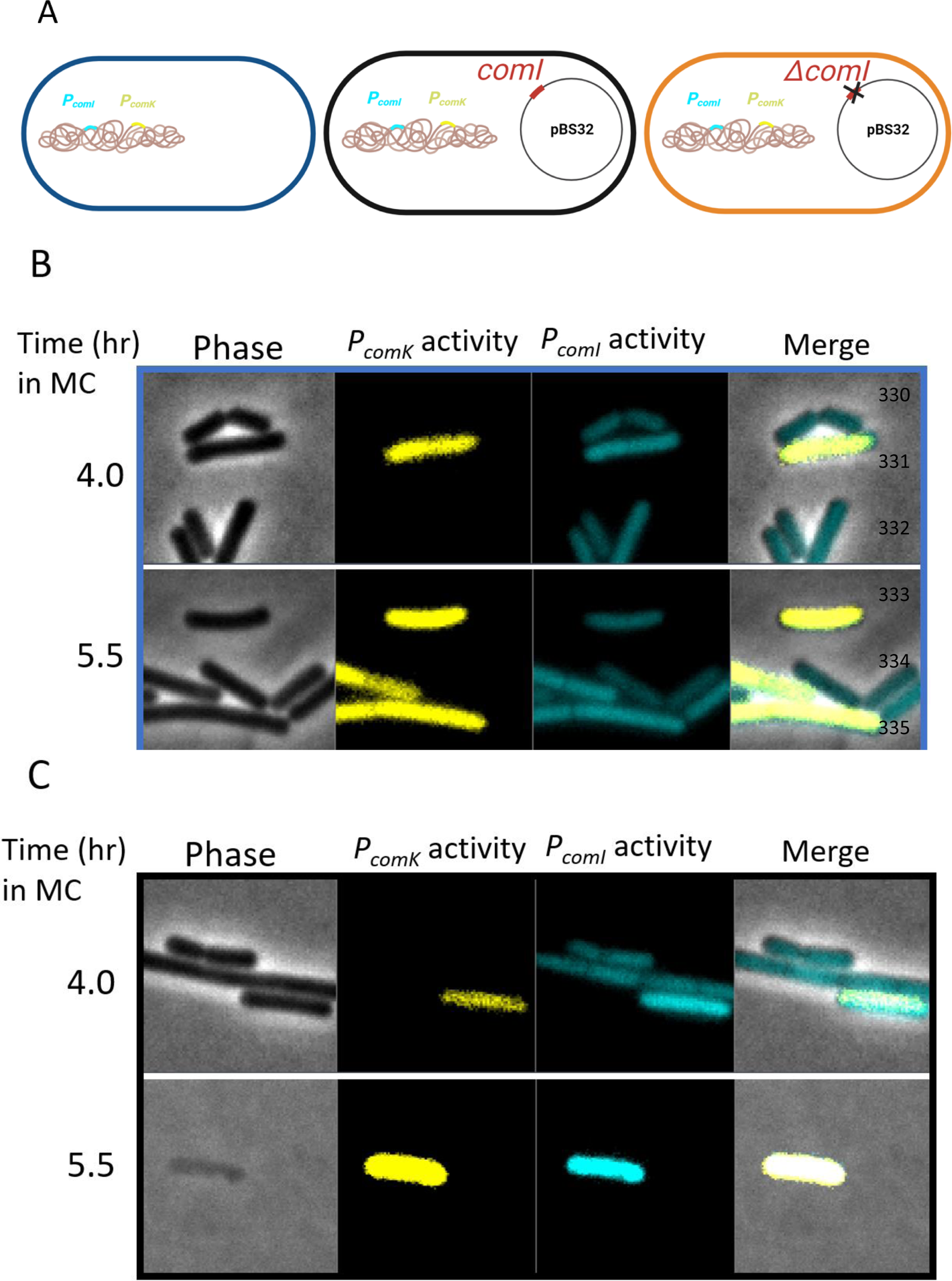
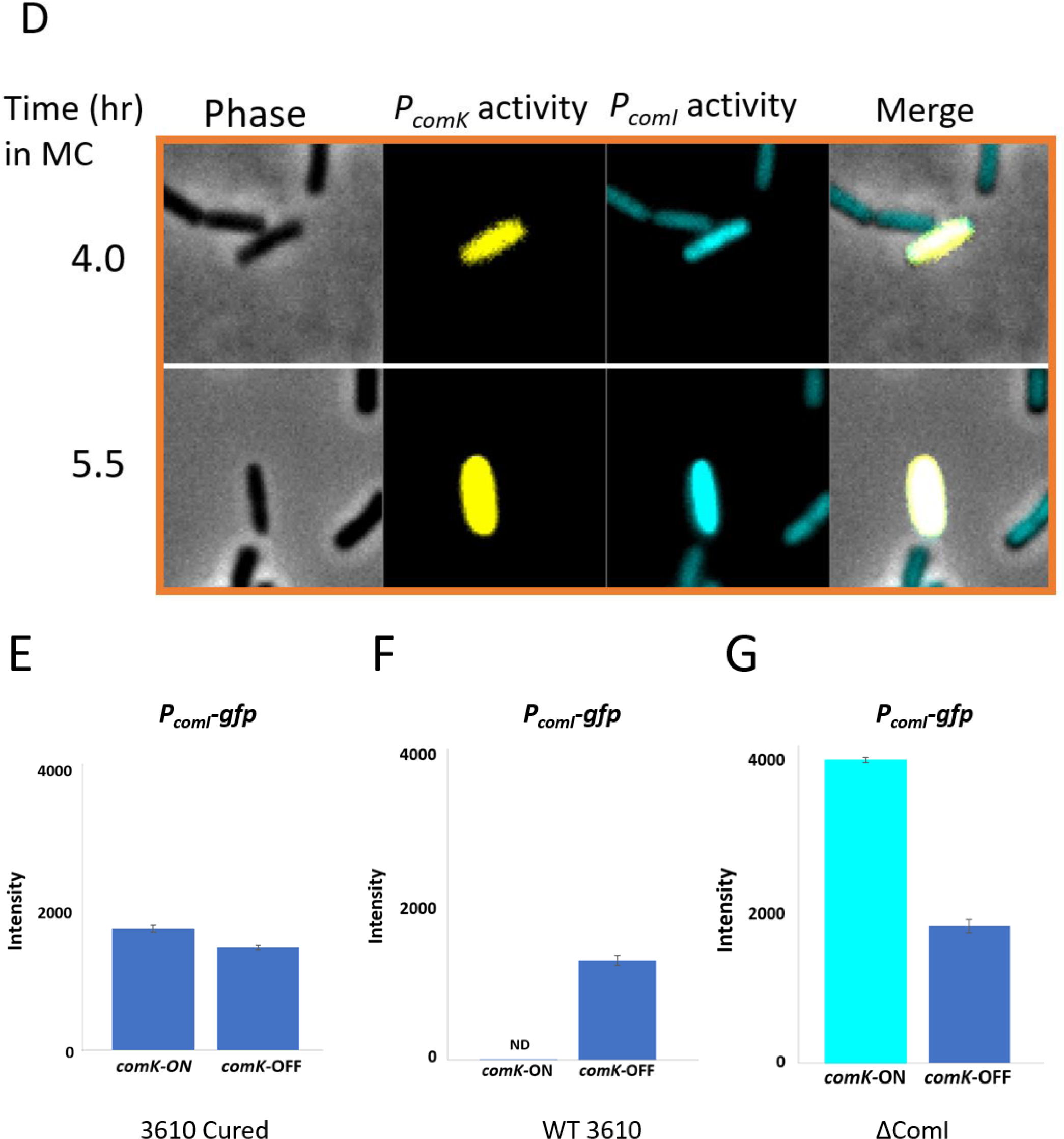
*comK*-ON cells have increased transcription from the native *comI* promoter. A) Schematic genotypes of cells used in this experiment. The colors (dark blue, black, and orange) respectively align with the three series of micrographs in B-D. B) Micrographs of 3610 cured cells C) Micrographs of WT 3610 cells D) Micrographs of cells in the *ΔcomI* background. All three micrographs include (from left to right): phase cells, cells with either *comI* (turquoise) or *comK* (yellow) reporters (both false-colored), and a merge of all three. E-G) Bar graphs depict the fluorescence intensity of GFP signal driven by the native *comI* promoter in cells that are *comK*-ON versus *comK*-OFF (n = 100 cells for each sample quantified). ND = Not Determined. No fluorescence intensity was determined for WT cells due to the infrequency of *comK*-ON cells in the population. The brighter blue color in this bar graph indicates that the intensity is higher. Strains used in this experiment include: bDS016, bDS017, and bDS022.

### ComI Selectively Permeabilizes the Competent Population Leading to Cell Death

Since inducible ComI permeabilizes the cells and *comI* reporter levels increase in competent cells, we asked if ComI, when expressed under its native promoter, permeabilizes the competent population over time. To assess competent cells for permeability, we integrated the *comK* transcriptional reporter in the WT 3610 genome and followed their uptake or PI. Cells were grown for four hours in competence conditions and then inoculated onto an agarose pad infused with PI. By tracking cells every five minutes via time-lapse microscopy, we determined that only the competent population had increases in PI signal above background (Figure 5A). These cells were phase-dark at 4 hours of growth in competence medium. However, by 5.5 hours, the cells were no longer phase-dark and appeared translucent, which indicates that they died (Figure 5A). All other cells remained viable in the population. Then, we conducted this same experiment in the Δ*comI* background. Here, we determined that over time, the competent population remained impermeable to the PI stain, and remained phase-dark at 5.5 hours of growth in competence medium (Figure 5B).

**Figure 5.**
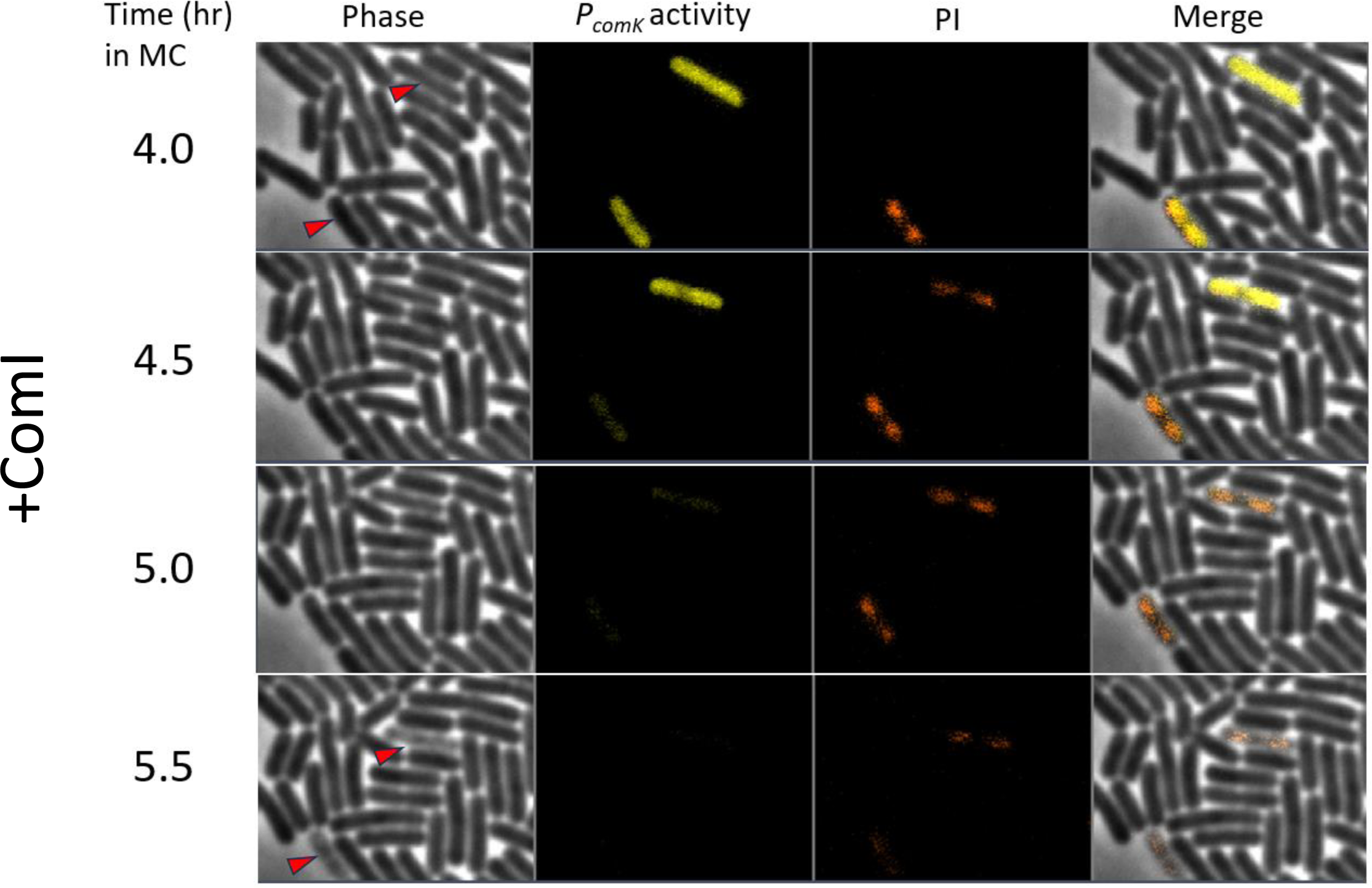
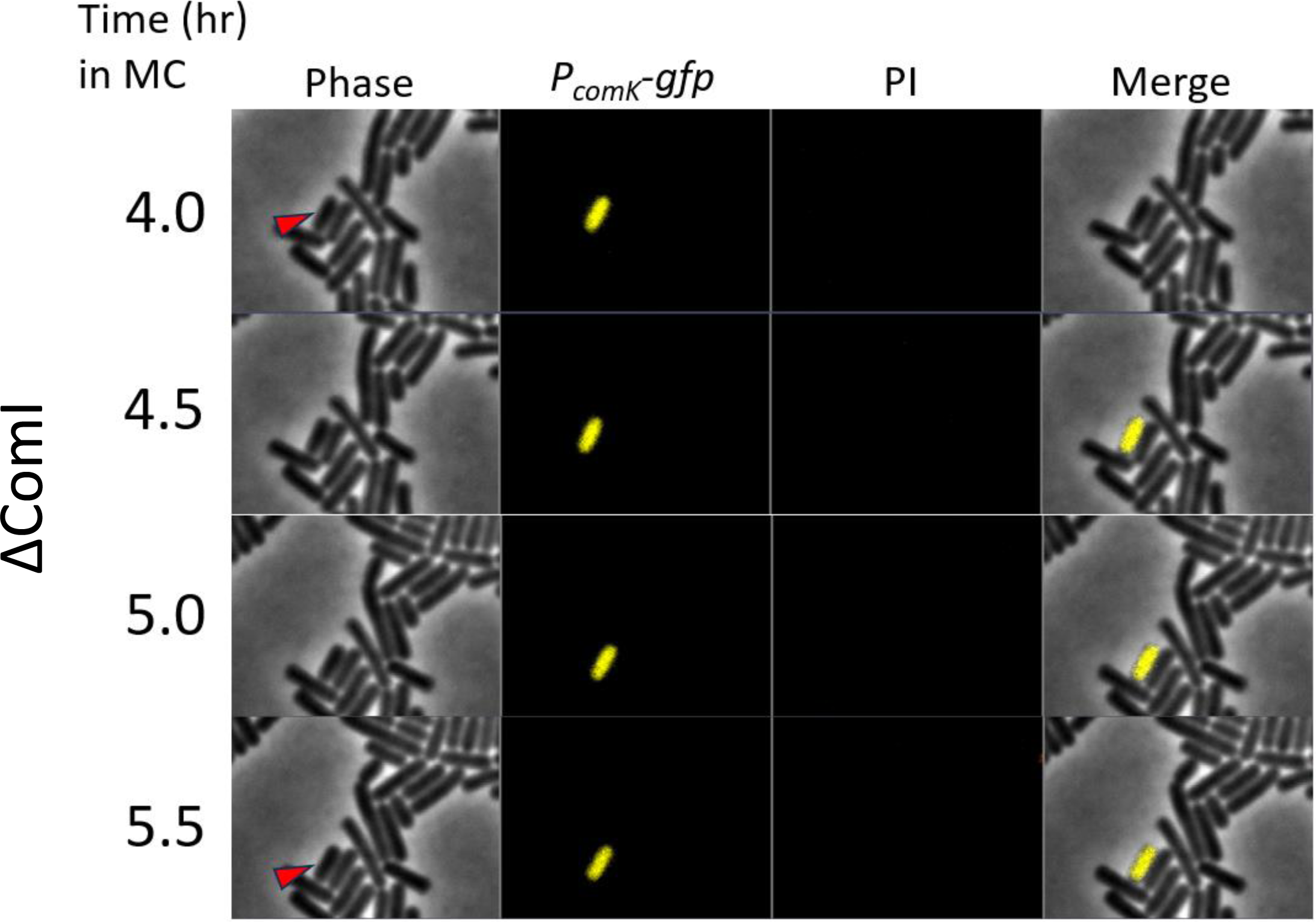
*comK*-ON cells become permeable dependent on the presence of ComI. A) Imaging of WT 3610 cells. B) Imaging of the *ΔcomI* background. Red arrows in the micrographs of each figure indicate the state of the cell (phase-dark or non-phase dark). The green color represents the signal gained from the activity of the *comK* promoter. Strains included in this figure: bDS090 and bDS089.

Based on these data, we propose that *B. subtilis* ComI selectively permeabilizes the competent subpopulation of cells leading to death (Figure 6A, 6B). This proposed mechanism of action is consistent with the function of a family of peptides that are homologous to ComI. Using Interpro to gain functional insights, we found that ComI is classified within the small toxic BsrE-like protein family (IPR031616) (22). This family comprises small single-pass transmembrane proteins that are toxic to gram-positive bacteria. Characterized members of this family are a part of type 1 toxin-antitoxin (TA) systems. Type 1 toxins primarily function by depolarization and/or permeabilization of the membrane in response to an environmental stressor, which aligns well with our functional data suggesting that ComI permeabilizes the cells in response to the competent state (23, 24).

**Figure 6.**
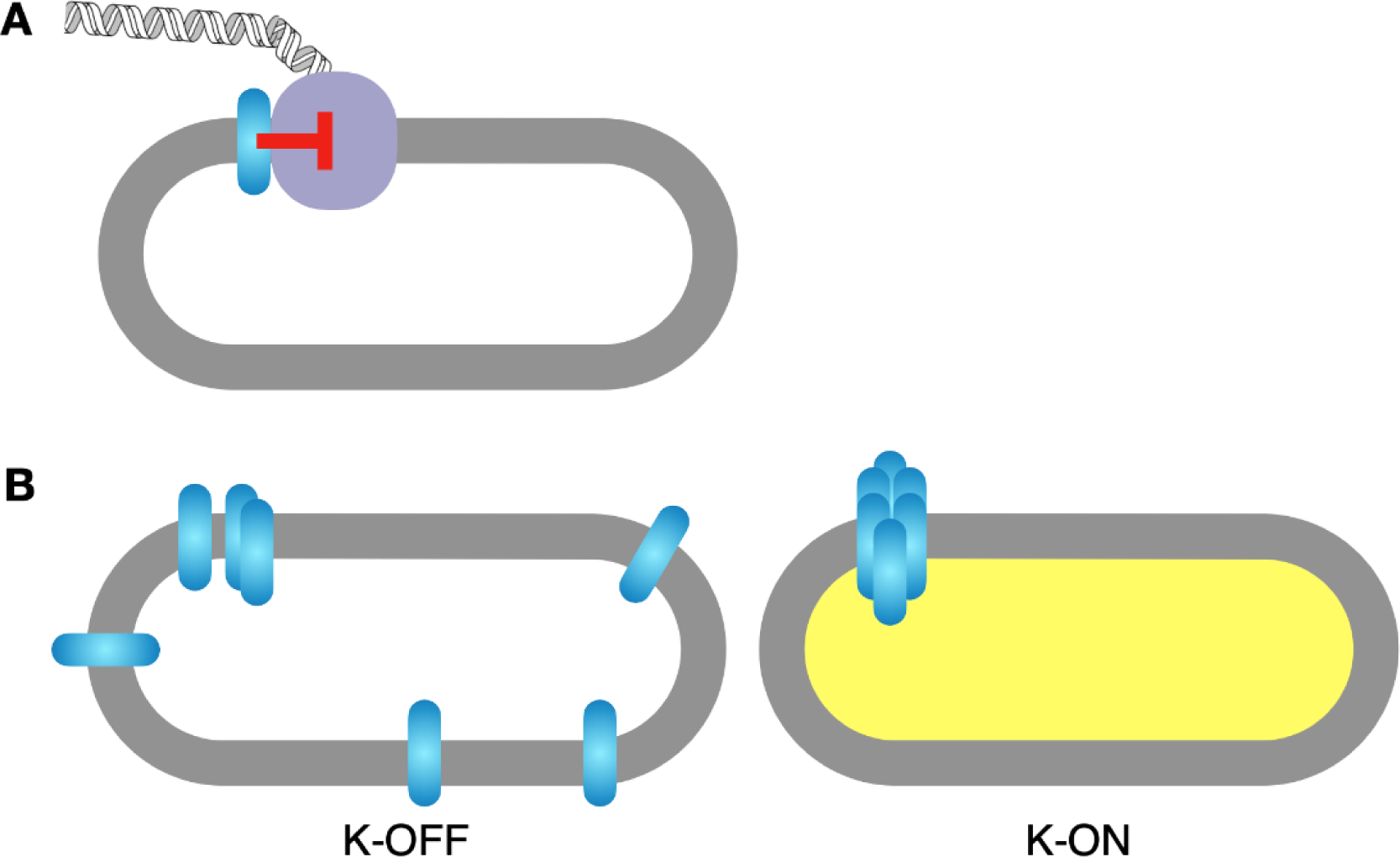
Model for ComI action. A) Schematic of previsouly proposed model in which ComI (cyan) interacts specifically with a protein or proteins from the transformation apparatus (purple) to inhibit DNA uptake. B) Selective permeabilization model proposed based on current data. The left cell depicts a non-competent cell surviving ComI in the membrane. The right cell depicts a competent cell being permeabilized by ComI. The altered physiology resulting from activation of the ComK regulon may potentiate ComI oligomerization.

## Discussion

The mechanism of action of the transformation inhibitor ComI has remained unstudied since its discovery a decade ago. One model suggests that ComI has the potential to inhibit transformation by directly interacting with competence proteins, although no interacting partner had been identified (10). However, the published observations of coordinated loss of transformation frequency followed closely by loss of viability in strains with inducible ComI, which provided alternative models for the potential ComI function. We found that inducible ComI permeabilizes cells at a critical concentration and does so independent of the competence machinery. These observations led us to postulate that ComI could be selectively permeabilizing and killing the competent subpopulation of cells. To test if this was happening at physiological levels of ComI, we ascertained ComI function under its endogenous promoter. In doing this, we discovered that the condition for native ComI expression required the presence of pBS32 as well as cell differentiation into the competent state. Finally, under these same conditions, time-lapse microscopy revealed that the competent population was permeabilized over time and that this was dependent on ComI. How differentiation into the competent state triggers permeabilization remains elusive.

Competent cells may be preferentially sensitive to ComI action due to ComK-driven altered physiology. In the 1960s, competent cells were shown to have different buoyant density from non-competent cells when they were first separated by centrifugation on a density gradient of Renografin-76 (25). The competent population was less dense than the noncompetent population, and therefore, the content of the cells was speculated to be different. Later gene expression studies supported this idea. Microarray and RNA-seq data indicate that genes associated with the cell wall are upregulated directly by ComK (5, 26). It is tempting to speculate that a difference in membrane lipid composition, fluidity, and cell wall components in *comK*-ON cells may provide the physical alterations necessary to sensitize cells to ComI activity.

Since ComI permeabilizes the cells, we speculated that a pore was forming, and therefore, ComI function could be dependent on self-interactions via oligomerization. This is consistent with the sharp concentration dependence of both the transformation and cell viability phenotypes (10). Other membrane-associated pore-forming peptides have been reported to oligomerize at critical concentrations to carry out their function. Specifically, phage holins accumulate harmlessly in the membrane until a signal triggers oligomerization and consequent permeabilization (27, 28). Similarly, some TA toxins can oligomerize and form pores (24, 29). It is possible that ComI is produced in non-competence conditions and remains in an inactive state until transition into the *comK*-ON state triggers oligomerization. Our reporter quantification suggests that ComI is transcribed at relatively low levels in the cell independent of competence and previously published REND-seq data supports this idea (Figure 4)(20, 21). Since, our data indicates that a critical concentration is necessary for inducible ComI to permeabilize and kill the cells, we speculate that oligomerization may be a prerequisite to ComI-mediated death (Figure 1E) (10).

It is puzzling why a plasmid, which is normally associated with horizontal gene transfer, encodes an inhibitor of competence with the propensity to kill competent cells. We have speculated two hypotheses. Firstly, this model draws parallels from the abortive infection systems for anti-phage defense. Abortive infection (Abi) is a commonly employed strategy by host cells to resist phages, ensuring that a phage epidemic does not propagate to nearby cells (30). These systems rely on sensing elements created by the phage genome and respond by activating their killing module, in this case ComI. Secondly, nearly half the genes on pBS32 encode for a putative prophage, and it has been speculated that the entire plasmid could be a prophage phagemid (16). Thus, this model could represent a phage defense system in response to DNA uptake. Considering that the competence state relies on proteins essential for the DNA damage response, competence can trigger SOS induction, and in turn, expose the ComI death mechanism to safeguard a population maintaining the prophage (31–36). Coupling inhibitor activity to the competent state can provide selective protection to the mobile element. This example would shed light on how a toxic peptide eliminates the competent cell population and ensures the persistence of the prophage genetic material. In either case, understanding inhibitors of competence has significant implications. Understanding inhibitor regulation and mechanism of action more deeply could lead to the development of genetic tools that make strains more or less transformable. Additionally, if organisms harbor competence inhibitors, removing them could yield significantly better transformation efficiencies.

The presence of inhibitors could explain why some species with preserved orthologous competence genes are incapable of transforming DNA. *Listeria monocytogenes* is a well-studied example, in which all the late competence genes are present and even upon excision of a prophage to allow expression of competence genes, transformation is still not observed (37). Other labs have reported that natural competence in *Staphylococcus aureus* cultures infrequently transform DNA (38, 39). Although experiments demonstrate that induction of *comK* results in transformable events while using plasmid DNA, chromosomal recombination events were very rare (40). Additionally, *Enterococci* are lagging other Gram-positive pathogens in molecular biology because of restrictions in DNA transfer (41). *Enterococci* contain elements necessary for natural competence, and it was suggested that they could play an alternative role in host colonization and immune invasion, but there has been no evidence for natural competence (42). It’s possible that non-competent bacteria containing orthologs of competence genes may contain an inhibitor, like *comI*, that integrates competence physiology with inhibitor activity to achieve transformation inhibition.

## Materials and Methods

### Strains and growth conditions

*B. subtilis* strains (Table 1) were grown in lysogeny broth (LB) broth (containing 10 g tryptone, 5 g yeast extract, and 5 g NaCl per liter) or on LB plates fortified with 1.5% Bacto agar at 37 °C. Modified competence (MC) medium (10X) was made with a solution containing 10.7 g K_2_HPO_4_, 5.2 g KH_2_PO_4_, 20 g dextrose anhydrous, 0.88 g sodium citrate dihydrate, 1 mL 1000X ferric ammonium citrate, 1 g casein hydrolysate, 2.2 g potassium glutamate monohydrate, and ddH_2_0 to a final volume of 100 mL. Competent cultures were grown in diluted 1X MC medium which was supplemented with 350 mM MgSO_4_. When appropriate, antibiotics were included at the following concentrations: 100 μg/mL spectinomycin, 1 μg/mL erythromycin plus 25 μg/mL lincomycin (*mls*), 5 μg/mL kanamycin, 5 μg/mL chloramphenicol, and 10 μg/mL tetracycline. Isopropyl-B-D-thiogalactopyranoside (IPTG) was added to cultures when appropriate.

**Table 1:** Strain List.

| Strain | Background | Genotype | Reference |
| --- | --- | --- | --- |
| bBB584 | PY79 | *amyE::P<sub>hyperspank</sub>-comI spec* | (10) |
| bDS001 | 3610 Cured | *amyE::P<sub>hyperspank</sub>-mCherry-comI spec* | (10) |
| bDS014 | 3610 Cured | *ΔcomK; amyE::P<sub>hyperspank</sub>-mCherry-comI spec* | This work |
| bLH080 | 3610 | WT | (10) |
| bBB592 | 3610 | *ΔcomI* | (10) |
| bJF176 | 3610 | ComI Q12L | (10) |
| bBS108 | 3610 Cured | Cured | (10) |
| bDS005 | 3610 Cured | *ytol::P<sub>comI</sub>-comI tet* | This work |
| bDS004 | 3610 | *ΔcomI; ytol::P<sub>comI</sub>-comI tet* | This work |
| bDS003 | 3610 | ComI Q12L *ytol::P<sub>comI</sub>-comI tet* | This work |
| bDS033 | 3610 | *ΔQIII; ytol::P<sub>comI</sub>-gfp tet* | This work |
| bDS092 | 3610 | *ΔQIII; ytol::P<sub>comI</sub>-gfp tet amyE::P<sub>hyperspank</sub>-zpbW mls* | This work |
| bDS016 | 3610 Cured | *ytol::P<sub>comI</sub>-gfp tet; lacA::P<sub>comK</sub>-mKate mls* | This work |
| bDS017 | 3610 | *ytol::P<sub>comI</sub>-gfp tet; lacA::P<sub>comK</sub>-mKate mls* | This work |
| bDS022 | 3610 | *ΔcomI; ytol::P<sub>comI</sub>-gfp; tet lacA::P<sub>comK</sub>-mKate mls* | This work |
| bDS089 | 3610 | *ΔcomI; lacA::P<sub>comK</sub>-gfp mls* | This work |
| bDS090 | 3610 | *lacA::P<sub>comK</sub>-gfp mls* | This work |
| bLH050 | 3610 Cured | *amyE::P<sub>hyperspank</sub>-empty* | This work |

For the transformation frequency assay, 1 mL LB cultures were inoculated and grown for 2 hours. Cultures were normalized to an optical density at 600 nm (OD_600_) of 0.05 in MC medium and grown rolling at 37 °C for 4.5 hours. Two hundred microliters of the culture were transferred into a 13 mm tube and 200 ng of genomic DNA (*yhdG::kan*) was added. The cultures incubated for an additional 2 hours at 250 rpm at 37 °C and were plated onto LB plates containing kanamycin to determine the number of transformants, and onto LB plates to determine the total viability. The transformation frequencies were determined by calculating the ratio of transforming units to colony forming units.

### Preparation of glass coverslips and agar pads for microscopy

All cover glasses used in microscopy experiments were pre-cleaned prior to use. 22 mm × 22 mm #1.5 borosilicate coverslips (DOT Scientific) were placed in a Wash-N-Dry coverslip rack (Sigma) and submerged in ∼80 mL of 1 M NaOH in a 100-mL glass beaker. The beaker was then placed into an ultrasonic cleaning bath (frequency = 40 kHz, power = 120 W) and sonicated for 30 minutes to remove the thin grease layer present on the coverslips. The coverslip rack was submerged into a fresh 250-mL glass beaker filled with ddH_2_O, the ddH_2_O was removed, and then the coverslips were washed 3× with 250 mL of ddH_2_O. The coverslips were then either air-dried overnight or dried immediately with compressed air.

For the preparation of agar pads, pre-cleaned borosilicate glass microscope slides were first rinsed free of detritus using ddH_2_O and were then either air-dried overnight or dried immediately with compressed air. SecureSeal 0.24 mm thick double sided adhesive sheets were cut into square pieces (approximately 0.5 inches in length and height), and a hole puncher was used to make a hole in the center of each square for placement of the agar pad. One side of the adhesive was removed, and this was centered on the borosilicate glass microscope slide. Then, the area of the glass slide that contained the hole was treated with 0.2 % chitosan to immobilize the agar pad. For the agar pad to be produced, approximately 15 minutes prior to imaging, conditioned MC medium from the cultures grown to competence via the transformation efficiency assay protocol was heated to 50°C and added in the ratio of 1:1 to 90°C 2% (wt/vol) molten LE agarose (SeaKem), and then vortexed to make molten 1% (wt/vol) agarose in 0.5× conditioned MC medium (pre-conditioned media was used when time-lapse microscopy was conducted, otherwise, molten LE agarose was mixed 1:1 with 1X PBS). Twenty microliters of this mixture were applied to the surface of the glass slide that was treated with chitosan, and immediately, another pre-cleaned glass slide was used to compress the agar pad down to the height of the adhesive sheet and was held in place for approximately 30 seconds so the agar could solidify. Once solidified, the slide was gently removed. Then, the second side of the SecureSeal adhesive sheet was removed to expose the adhesive and 1 μL of culture was added to the agarose pad. Finally, pre-treated coverslips were placed over the agar pad and adhesive and contact between the two sealed the agar pad in place. This specific setup allows for the agar pad to stick to the bottom slide and easily slide out from the top slide, leaving an unmarred and flat surface for imaging.

### Microscopy

All single time point imaging was performed using a Zeiss Axio Imager.Z2 upright epifluorescence microscope equipped with a Zeiss Plan-Apochromat 100×/1.4 Oil PH3 objective lens and a Teledyne Photometrics Prime 95B sCMOS camera. For fluorescence microscopy using the *comI* and *comK* transcriptional fusions, cells were grown in 1 mL of LB broth for 2 hours at 37 °C. Cells were centrifuged at 10,000 x g for 5 minutes, resuspended in approximately 15 % residual media, normalized to an optical density of 0.05 at a wavelength of 600 nm, and were thereafter grown at 37 °C until competence. For each time point, a total of 200 μL of culture was removed from the culture, harvested by centrifugation (8,000 x g for 5 minutes), and resuspended in 10 μL of phosphate-buffered saline (PBS) with a pH adjusted to 7.4. One μL of the suspension was spotted and immobilized onto an agarose pad. Cells were exposed to light from a Zeiss Colibri 580 nm LED module with Zeiss filter set 14 alexa fluor 546 for 2000 ms at 20% LED power to observe *PcomK-mKate*, while *PcomI-gfp* was observed using light from a Zeiss Colibri 475 nm LED module with Zeiss filter set 38 HE for 100 ms at 20% LED power. Cell bodies were imaged using phase-contrast microscopy with 30 ms exposures using a TL halogen lamp with a light intensity of 3.40 V.

For fluorescence microscopy of the mCherry-ComI translational fusion, cells were grown in 1 mL of LB broth for 3 hours, and then induced with 70 μM IPTG for 1.5 hours. Cells were harvested by centrifuging 1 mL of culture followed by resuspension in 50 μL of PBS (pH = 7.4). One microliter of the suspension was spotted and immobilized onto an agarose pad. The exposure time for mCherry-ComI was 2000 ms.

For fluorescence microscopy of inducible ComI strains that were stained with PI, cells were grown in LB for 3 hours, then incubated with IPTG for 1.5 hours before they were harvested and deposited onto 2% agarose pads infused with 300ng/μL propidium iodide. Cells were exposed to light from a Zeiss Colibri 580 nm LED module with Zeiss filter set 14 alexa fluor 546 for 250 ms at 20% LED power to observe PI stain.

### Time-lapse microscopy imaging

Cells were grown in 1 mL of LB broth for 2 hours at 37 °C. Cells were centrifuged at 10,000 x g for 5 minutes, resuspended in approximately 15 % residual media, normalized to an optical density of 0.05 at a wavelength of 600 nm, and were thereafter grown in 1X MC at 37 °C for 4 hours. Then, cells were centrifuged at 10,000 x g for 5 minutes and resuspended in 15 % residual supernatant (the remaining pre-conditioned supernatant was used to make the agarose pad; see instructions for preparation of agarose pad above), followed by deposition onto the pre-conditioned agarose pad. Before imaging began, a PECON Heater S objective heater calibrated with thermocouple was set to 37 °C and was placed around the objective lens. The prepared sample slide was placed on the heated objective. For imaging, a time-series was conducted that took 1 image in all three channels every 5 minutes for 2 hours. The focus strategy utilized was definite focus (referenced to phase channel) and was set to re-focus during the acquisition of each image. Cells were imaged using a Zeiss Axio Observer.Z1 inverted epifluorescence microscope equipped with a Zeiss Plan-Apochromat 100×/1.4 Oil PH3 M27 objective lens and a Teledyne Photometrics CoolSNAP HQ2 CCD camera. Cells were exposed to light from a Zeiss Colibri 470 nm LED module with Zeiss filter set 14 alexa fluor 546 for 1000 ms at 20% LED power to observe *PcomK-gfp*, while propidium iodide stain was observed using light from a Zeiss Colibri 590 nm LED module with Zeiss filter set 38 HE for 250 ms at 20% LED power. Cell bodies were exposed to light for 30 ms from a TL halogen lamp with an intensity of 6.3 V and were imaged using phase-contrast microscopy.

## References

1. Blokesch M. 2016. Natural competence for transformation. Current Biology 26(21):R1126–R1130.

2. Redfield RJ. 1993. Genes for breakfast: The have-your-cake and-eat-lt-too of bacterial transformation. Journal of Heredity 84(5):400–4.

3. Johnston C, Martin B, Fichant G, Polard P, Claverys JP. 2014. Bacterial transformation: Distribution, shared mechanisms and divergent control. Nat Rev Microbiol 12(3):181–96.

4. Winter M, Buckling A, Harms K, Johnsen PJ, Vos M. 2021. Antimicrobial resistance acquisition via natural transformation: context is everything. Curr Opin Microbiol 64:133–138.

5. Berka RM, Hahn J, Albano M, Draskovic I, Persuh M, Cui X, Sloma A, Widner W, Dubnau D. 2002. Microarray analysis of the Bacillus subtilis K-state: Genome-wide expression changes dependent on ComK. Mol Microbiol 43(5):1331–45.

6. Smith MCM. 1991. Molecular biological methods for bacillus. FEBS Lett 287.

7. Nye TM, Schroeder JW, Kearns DB, Simmons LA. 2017. Complete genome sequence of undomesticated Bacillus subtilis strain NCIB 3610. Genome Announc 5.

8. Zeigler DR, Prágai Z, Rodriguez S, Chevreux B, Muffler A, Albert T, Bai R, Wyss M, Perkins JB. 2008. The origins of 168, W23, and other Bacillus subtilis legacy strains. J Bacteriol 190.

9. Nijland R, Burgess JG, Errington J, Veening JW. 2010. Transformation of environmental Bacillus subtilis isolates by transiently inducing genetic competence. PLoS One 5.

10. Konkol MA, Blair KM, Kearns DB. 2013. Plasmid-encoded comi inhibits competence in the ancestral 3610 strain of Bacillus subtilis. J Bacteriol 195.

11. Earl AM, Losick R, Kolter R. 2007. Bacillus subtilis genome diversity. J Bacteriol 189.

12. Haijema BJ, Hahn J, Haynes J, Dubnau D. 2001. A ComGA-dependent checkpoint limits growth during the escape from competence. Mol Microbiol 40.

13. Provvedi R, Dubnau D. 1999. ComEA is a DNA receptor for transformation of competent Bacillus subtilis. Mol Microbiol 31.

14. Draskovic I, Dubnau D. 2005. Biogenesis of a putative channel protein, ComEC, required for DNA uptake: Membrane topology, oligomerization and formation of disulphide bonds. Mol Microbiol 55.

15. Crowley LC, Scott AP, Marfell BJ, Boughaba JA, Chojnowski G, Waterhouse NJ. 2016. Measuring cell death by propidium iodide uptake and flow cytometry. Cold Spring Harb Protoc 2016.

16. Burton AT, Kearns DB. 2020. The large pBS32/pLS32 plasmid of ancestral bacillus subtilis. J Bacteriol 202(18):e00290–20.

17. Turgay K, Hamoen LW, Venema G, Dubnau D. 1997. Biochemical characterization of a molecular switch involving the heat shock protein ClpC, which controls the activity of ComK, the competence transcription factor of Bacillus subtilis. Genes Dev 11.

18. Maamar H, Dubnau D. 2005. Bistability in the Bacillus subtilis K-state (competence) system requires a positive feedback loop. Mol Microbiol 56.

19. Smits WK, Eschevins CC, Susanna KA, Bron S, Kuipers OP, Hamoen LW. 2005. Stripping Bacillus: ComK auto-stimulation is responsible for the bistable response in competence development. Mol Microbiol 56.

20. DeLoughery A, Lalanne JB, Losick R, Li GW. 2018. Maturation of polycistronic mRNAs by the endoribonuclease RNase Y and its associated Y-complex in Bacillus subtilis. Proc Natl Acad Sci U S A 115.

21. Burton AT, DeLoughery A, Li GW, Kearns DB. 2019. Transcriptional regulation and mechanism of sigN (ZpdN), a pBS32-encoded sigma factor in bacillus subtilis. mBio 10.

22. Paysan-Lafosse T, Blum M, Chuguransky S, Grego T, Pinto BL, Salazar GA, Bileschi ML, Bork P, Bridge A, Colwell L, Gough J, Haft DH, Letunić I, Marchler-Bauer A, Mi H, Natale DA, Orengo CA, Pandurangan AP, Rivoire C, Sigrist CJA, Sillitoe I, Thanki N, Thomas PD, Tosatto SCE, Wu CH, Bateman A. 2023. InterPro in 2022. Nucleic Acids Res 51.

23. Nonin-Lecomte S, Fermon L, Felden B, Pinel-Marie ML. 2021. Bacterial type I toxins: Folding and membrane interactions. Toxins (Basel) 13(7):490.

24. Brielle R, Pinel-Marie ML, Felden B. 2016. Linking bacterial type I toxins with their actions. Curr Opin Microbiol 30:114–121.

25. Hadden C, Nester EW. 1968. Purification of competent cells in the Bacillus subtilis transformation system. J Bacteriol 95.

26. Boonstra M, Schaffer M, Sousa J, Morawska L, Holsappel S, Hildebrandt P, Sappa PK, Rath H, de Jong A, Lalk M, Mäder U, Völker U, Kuipers OP. 2020. Analyses of competent and non-competent subpopulations of Bacillus subtilis reveal yhfW, yhxC and ncRNAs as novel players in competence. Environ Microbiol 22.

27. Shi Y, Yan Y, Ji W, Du B, Meng X, Wang H, Sun J. 2012. Characterization and determination of holin protein of Streptococcus suis bacteriophage SMP in heterologous host. Virol J 9.

28. Saier MH, Reddy BL. 2015. Holins in bacteria, eukaryotes, and archaea: Multifunctional xenologues with potential biotechnological and biomedical applications. J Bacteriol 197(1):7–17.

29. Thompson MK, Nocedal I, Culviner PH, Zhang T, Gozzi KR, Laub MT. 2022. Escherichia coli SymE is a DNA-binding protein that can condense the nucleoid. Mol Microbiol 117.

30. Lopatina A, Tal N, Sorek R. 2020. Abortive Infection: Bacterial Suicide as an Antiviral Immune Strategy. Annu Rev Virol 7(1):371–384.

31. Mcveigh RR, Yasbin RE. 1996. Phenotypic differentiation of “smart” versus “naive” bacteriophages of Bacillus subtilis. J Bacteriol 178.

32. Dorer MS, Fero J, Salama NR. 2010. DNA damage triggers genetic exchange in Helicobacter pylori. PLoS Pathog 6.

33. Yasbin RE, Wilson GA, Young FE. 1975. Transformation and transfection in lysogenic strains of Bacillus subtilis: evidence for selective induction of prophage in competent cells. J Bacteriol 121.

34. Garro AJ, Law MF. 1974. Relationship between lysogeny, spontaneous induction, and transformation efficiencies in Bacillus subtilis. J Bacteriol 120.

35. Okamoto K, Mudd JA, Marmur J. 1968. Conversion of Bacillius subtilis DNA to phage DNA following mitomycin C induction. J Mol Biol 34.

36. Goranov AI, Kuester-Schoeck E, Wang JD, Grossman AD. 2006. Characterization of the global transcriptional responses to different types of DNA damage and disruption of replication in Bacillus subtilis. J Bacteriol 188.

37. Rabinovich L, Sigal N, Borovok I, Nir-Paz R, Herskovits AA. 2012. Prophage excision activates listeria competence genes that promote phagosomal escape and virulence. Cell 150.

38. Morikawa K, Takemura AJ, Inose Y, Tsai M, Nguyen Thi LT, Ohta T, Msadek T. 2012. Expression of a Cryptic Secondary Sigma Factor Gene Unveils Natural Competence for DNA Transformation in Staphylococcus aureus. PLoS Pathog 8.

39. Thi LTN, Romero VM, Morikawa K. 2016. Cell wall-Affecting antibiotics modulate natural transformation in SigH-expressing Staphylococcus aureus. Journal of Antibiotics 69(6):464–6.

40. Cordero M, García-Fernández J, Acosta IC, Yepes A, Avendano-Ortiz J, Lisowski C, Oesterreicht B, Ohlsen K, Lopez-Collazo E, Förstner KU, Eulalio A, Lopez D. 2022. The induction of natural competence adapts staphylococcal metabolism to infection. Nat Commun 13.

41. Krause AL, Stinear TP, Monk IR. 2022. Barriers to genetic manipulation of Enterococci: Current Approaches and Future Directions. FEMS Microbiol Rev 46(6):fuac036.

42. Abril AG, Quintela-Baluja M, Villa TG, Calo-Mata P, Barros-Velázquez J, Carrera M. 2022. Proteomic Characterization of Virulence Factors and Related Proteins in Enterococcus Strains from Dairy and Fermented Food Products. Int J Mol Sci 23.

43. Gibson DG, Young L, Chuang RY, Venter JC, Hutchison CA, Smith HO. 2009. Enzymatic assembly of DNA molecules up to several hundred kilobases. Nat Methods 6.

